# A physiological drop in pH decreases mitochondrial respiration, and AMPK and Akt signalling, in L6 myocytes

**DOI:** 10.1101/330662

**Authors:** Amanda J Genders, Sheree D Martin, Sean L McGee, David J Bishop

## Abstract

Exercise stimulates mitochondrial biogenesis and increases mitochondrial respiratory function and content. However, during high-intensity exercise muscle pH can decrease below pH 6.8 with a concomitant increase in lactate concentration. This drop in muscle pH is associated with reduced exercise-induced mitochondrial biogenesis, whilst increased lactate may act as a signaling molecule to affect mitochondrial biogenesis. Therefore, in this study we wished to determine the impact of altering pH and lactate concentration in L6 myotubes on genes and proteins known to be involved in mitochondrial biogenesis. We also examined mitochondrial respiration in response to these perturbations. Differentiated L6 myotubes were exposed to normal (pH 7.5), low (pH 7.0) or high pH (pH 8.0) media with and without 20 mM sodium L-lactate for 1 and 6 h. Low pH and 20 mM Sodium L-Lactate resulted in decreased Akt (Ser473) and AMPK (T172) phosphorylation at 1 h compared to controls, whilst at 6 h the nuclear localisation of HDAC5 was decreased. When the pH was increased both Akt (Ser473) and AMPK (T172) phosphorylation was increased at 1 h. Overall increased lactate decreased the nuclear content of HDAC5 at 6 h. Exposure to both high and low pH media significantly decreased basal mitochondrial respiration, ATP turnover, and maximum mitochondrial respiratory capacity. These data indicate that muscle pH affects several metabolic signalling pathways, including those required for mitochondrial function.

## Introduction

Exercise stimulates mitochondrial biogenesis, leading to an increase in mitochondrial content and respiratory function, and this has been attributed to the cumulative effects of each single exercise session [1-6]. This process is initiated in response to multiple perturbations of cellular homeostasis (e.g., increases in the ADP/ATP ratio) [7], which are followed by the activation of kinases such as AMP-activated protein kinase (AMPK), Ca^2+^/calmodulin-dependent protein kinase (CaMK), and mitogen-activated protein kinase (p38 MAPK) [2, 8]. These signalling pathways have all been reported to activate and/or increase the expression of proliferator-activated receptor γ coactivator 1 α (PGC-1α), a transcriptional coactivator that interacts with transcription factors, such as nuclear respiratory factor 1 (NRF-1), myocyte-enhancing factor-2 (MEF2), and mitochondrial transcription factor A (Tfam) [9], to up-regulate the mRNA and protein expression of mitochondrial genes and proteins [2].

One of the many cellular perturbations with exercise is an increase in muscle lactate concentration [10]. Blood lactate concentrations of approximately 15 to 25 mmol.L^-1^ may be observed immediately post high-intensity exercise [11, 12]. In the only study to date, genes implicated in exercise-induced mitochondrial biogenesis (e.g. NRF-2, COX-IV and PGC- 1α) were increased up to two fold in L6 myotubes that had been incubated with 20 mM of sodium lactate for six hours [13]. Thus, it was suggested that lactate may act as a signaling molecule to affect mitochondrial biogenesis in response to exercise [13]. The authors further hypothesized that the mechanism may be related to signaling through CaMKII and p38 MAPK via increased production of reactive oxygen species (ROS), although this was not directly measured [13]. However, lactate did increase hydrogen peroxide production four fold, and it also upregulated genes known to be responsive to ROS and calcium. The authors concluded that the lactate signaling cascade involves ROS production and converges on transcription factors affecting mitochondrial biogenesis. However, these results have not been replicated and many of the changes were small (< 1.4 fold).

During high-intensity exercise, lactate accumulation does not occur in isolation and is associated with an increase in the hydrogen ion concentration; this results in a decrease in muscle pH to values as low as pH 6.8 in the soleus and 6.6 in the EDL of rats [14], with a similar decrease in the vastus lateralis muscle of active women [15]. This decrease in pH is sufficient to have an effect on metabolism [16] and to alter the expression and/or activity of some proteins (e.g., basal insulin receptor substrate-1 (IRS-1) associated phosphatidylinositol 3-kinase (PI3-K), ubiquitin, and protease subunit mRNA [17-19]). A lower muscle pH in humans has also been associated with a reduced exercise-induced expression of genes known to be involved in mitochondrial biogenesis (e.g., PGC-1α) [20]. In rats, administration of ammonium chloride, resulting in a lowering of blood pH from 7.38 to 7.16, decreased MAPK phosphorylation in the kidney [21]. In a study with HeLa cells, the lowering of intracellular pH (via the manipulation of sodium bicarbonate levels) decreased histone acetylation and affected the expression of many genes including those in the MAPK signalling pathway [22]. To date, however, no study (with the exception of an abstract by Perez-Schindler et al [23]) has investigated the effects of manipulating pH on cell signaling pathways associated with mitochondrial biogenesis in myocytes.

There is therefore, some evidence to suggest two cellular perturbations that occur with high-intensity exercise (increased lactate concentration and decreased muscle pH) may act on genes and proteins implicated in exercise-induced mitochondrial biogenesis. However, although some of these factors have been studied independently in muscle cell culture, no study has looked at these two manipulations together and no study has examined in detail genes and proteins known to be involved in mitochondrial biogenesis. The aim of this study was to determine the impact of altering pH (by changing bicarbonate concentration), with and without an increase in media lactate concentration, in L6 myotubes. In particular, we examined changes in genes and proteins involved in the regulation of exercise-induced mitochondrial biogenesis, as well as the effect of these two cellular perturbations on mitochondrial respiration. To enable comparison with previous literature, we have performed experiments in both low and high glucose containing media. It was hypothesized that changes in both pH and lactate concentration would affect the expression of genes and proteins associated with mitochondrial biogenesis in L6 cells.

## Methods

### Cell culture

L6 myoblasts (American Tissue Culture Collection) were cultured in Minimum Essential Media (MEM) α (5.5 mmol/L glucose, 10% foetal bovine serum, 1% antibiotic/antimycotic)(low glucose) or Dulbecco’s Modified Essential Media (DMEM) (25 mmol.L^-1^ glucose, 10% foetal bovine serum) (high glucose) (Thermo Fisher Scientific, Melbourne, Australia) and seeded into 6 or 96 well plates for experimental measurements. Two different glucose concentrations were used in order to compare with previously-published data [13]. Cells were differentiated into myotubes by changing the serum to 2% horse serum (Thermo Fisher Scientific, Melbourne, Australia). The differentiation medium was replaced every 48 h. The identity of cells was assessed by surveying mRNA expression of myogenic differentiation-1 (MyoD) and myosin heavy chain-2 (Myh2) myocyte genes with qPCR and differentiation was confirmed by light microscopy. Mycoplasma contamination tests were not carried out. Differentiated L6 myocytes (5 to 6 days post-differentiation) were treated with normal, low, or high pH media, with and without the addition of 20 mM sodium L-lactate as used in a previous study [13], for zero, one or six hours. The incubation values of 20 mM sodium lactate and a pH of approximately 6.8 were chosen as similar values have been observed in human skeletal muscle after physical activity [20, 24, 25]. This gave the following groups: Normal pH, Normal pH + 20 mM Sodium Lactate, High pH, High pH + 20 mM Sodium Lactate, Low pH, and Low pH + 20 mM Sodium Lactate. The pH of the cell culture media was altered by increasing or decreasing the sodium bicarbonate concentration resulting in a pH of 8.04 ± 0.02 (high) and 6.97 ± 0.03 (low), respectively, as well as a normal pH of 7.57 ± 0.03, after incubation at 37°C and 5% CO_2_ for one hour. We also verified that there were concomitant changes in intracellular pH (Figure 1, described below). Cell viability was measured using trypan blue staining and a commercial LDH cytotoxicity assay (Thermo Fisher Scientific, Melbourne, Australia). Glucose and lactate concentrations in the media were measured with a Glucose Lactate analyser (YSI 2300 STAT Plus, John Morris Scientific, Melbourne, Australia).

**Figure 1:**
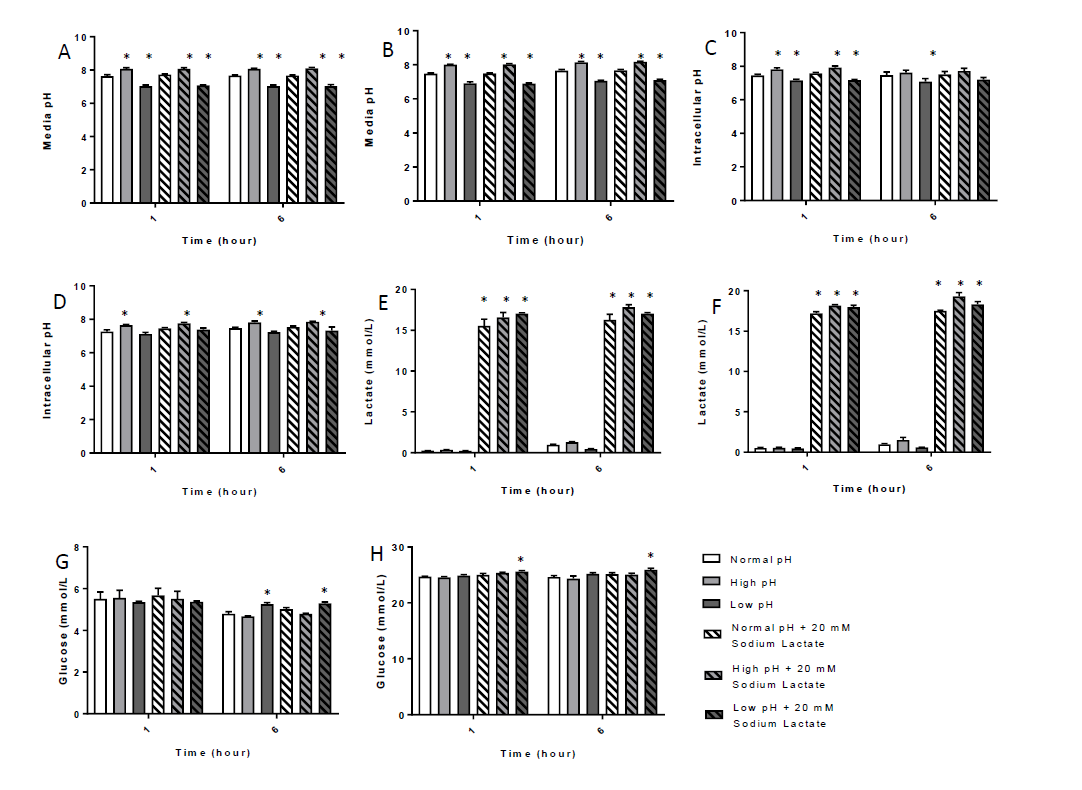
Incubation of L6 myocytes in low or high glucose media with normal, high, or low pH and +/- 20 mM Sodium Lactate. A. Low glucose media pH B. High glucose media pH C. Low glucose intracellular pH D. High glucose intracellular pH E. Low glucose media lactate F. High glucose media lactate G. Low glucose media glucose H. High glucose media glucose * significantly different from Normal pH group. P ≤ 0.05. Values are means ± SEM. Samples are from 4 to 5 individual experiments.

### Intracellular pH measurement

Intracellular pH was measured using 5-(-6)-carboxy SNARF®-1, acetoxymethyl ester, acetate (Thermo Fisher Scientific, Melbourne, Australia) in fully-differentiated L6 myocytes. The method was adapted from Behbahan et al [26]. Briefly 10 μM SNARF-1 with Pluronic F127 (Thermo Fisher Scientific, Melbourne, Australia) diluted in 1x EBSS with 1 g/L glucose and 24 mM NaHCO3 was loaded into the cells for 50 min at 37°C. Cells were then washed with PBS to remove excess dye and incubated with the different pH medias (pH 7.0, 7.5 and 8.1), with and without 20 mM sodium lactate, for 1 or 6 h. To establish a calibration curve, individual wells were incubated with calibration buffer (135 mM KCl, 2 mM K_2_HPO4, 20 mM HEPES, 1.2 mM CaCl_2_, 0.8 mM MgSO4) at the following pH: 6.0, 6.5, 7.0, 8.0 and 8.5 with 10 μM nigericin for 5 minutes at 37°C. Fluorescence was read on a plate reader with excitation at 530 nm and emission at 580 nm and 640 nm. Intracellular pH was calculated ratiometrically using a sigmoidal 4-parameter curve fit (SoftMax Pro 6.5.1).

### Western blotting

Total protein was extracted for analysis in ice cold lysis buffer (0.05M Tris pH 7.5, 1mM EDTA, 2mM EGTA, 10% glycerol, 1% Triton X-100, 1mM DTT) with the addition of a Protease and Phosphatase Inhibitor cocktail (Cell Signaling Technologies, Danvers, MA). Separation and purification of cytoplasmic and nuclear extracts from L6 myocytes was performed using a NE-PER Nuclear and Cytoplasmic Extraction kit (Thermo Fisher Scientific, Melbourne, Australia). Lysed samples were assayed for protein content and 5 to 10 μg protein was loaded onto TGX Stain-Free FastCast Acrylamide gels. Proteins were separated by electrophoresis and then transferred onto PVDF membrane using a standard protocol. Membranes were then blocked for 1 h at room temperature in TBST (TBS with 0.05% Tween 20 pH 7.4) with either 1% bovine serum albumin (BSA) or 5% skim milk powder. Membranes were then probed with the following primary antibodies overnight at 4°C at 1:1000 in TBST (all antibodies from Cell Signaling Technologies unless otherwise noted), phospho-p38 MAPK, total p38 MAPK, phospho-Ser473 Akt, total Akt, phospho-CaMKII, total CaMKII, phosphoT172AMPKα, total AMPKα, HDAC5, PGC-1α (Calbiochem – Merck Millipore, Darmstadt, Germany), Histone H3, LDH. Blots were then washed with TBST prior to incubation with the appropriate HRP-linked secondary antibody (Perkin Elmer) for 1 h at room temperature. Blots were developed using Clarity ECL and visualised using a ChemiDoc. All bands were quantified using ImageLab software (Bio-Rad Laboratories, Hercules, CA). All phosphorylated and individual protein expression was normalized to total protein. Purification of nuclear and cytosolic protein was confirmed by probing for Histone H3 and LDH. PGC-1α and HDAC5 abundance was determined in nuclear fractions.

### qPCR

RNA was extracted using TRIzol® Reagent (Thermo Fisher Scientific, Melbourne, Australia) as described in the manufacturer’s instructions. The purity of each sample was assessed from the A260/A280 absorption ratio using a BioPhotometer (Eppendorf AG, Hamburg, Germany). Total RNA concentration was also measured using the BioPhotometer. RNA integrity of a subset of the samples was measured using a Bio-Rad Experion microfluidic gel electrophoresis system (Bio-Rad, Hercules, CA) and determined from the RNA quality indicator (RQI). All samples were of a good quality (RQI 9.9 ± 0.01) and protein contamination was low (A260/A280 ratio was 2.03 ± 0.01). RNA was reverse transcribed to first strand cDNA from 1 μg of template RNA using a Thermocycler (Bio-Rad, Hercules, CA) and Bio-Rad iScript^TM^ RT Supermix (Bio-Rad, Hercules, CA) according to the kit instructions. qPCR for the following genes, MCT1 (Forward 5’-CGT TGA TGG ACC TCG TTG GA, Reverse 5’- CGA TGA TGA GGA TCA CGC CA), CD147 (Forward 5’- GGC GGG CAC CAT CGT AA, Reverse 5’- CCT TGC CAC CTC TCA TCC AG, NRF1 (Forward 5’-CTA CTC GTG TGG GAC AGC AA, Reverse 5’-AGC AGA CTC CAG GTC TTC CA), NRF2 (Forward 5’- AGT AGC GCA AAG GCA GCT AA, Reverse 5’- CCA TTG TTT CCT GTT CTG TTC CC), COXIV Forward 5’- GCA GCA GTG GCA GAA TGT TG, Reverse 5’-CGA AGG CAC ACC GAA GTA GA), Tfam (Forward 5’- AAT GTG GGG CGT GCT AAG AA, Reverse 5’- ACA GAT AAG GCT GAC AGG CG), PGC-1α (Forward 5’- ATA CAC AAC CGC AGT CGC AAC, Reverse 5’- GCA GTT CCA GAG AGT TCC ACA C), PGC-1α1 (Forward 5’-ATG GAG TGA CAT CGA GTG TGC Reverse 5’- GAG TCC ACC CAG AAA GCT GT), PGC-1α4 (Forward 5’-TCA CAC CAA ACC CAC AGA GA, Reverse 5’- CTG GAA GAT ATG GCA CAT), cytochrome c (Forward 5’- ATG GTC TGT TTG GGC GGA A, Reverse 5’- TCC CCA GGT GAT ACC TTT GTT C), p21 (Forward 5’- GCT GTC TTG CAC TCT GGT GT, Reverse 5’- AAT CGG CGC TTG GAG TGA TA), MyoD (Forward 5’- CAC TAC AGC GGC GAC TCA GA, Reverse 5’- TCA CTG TAG TAG GCG TC), and Myh2 (Forward 5’- GTG AAA ACT GAA GCA GGA GCG, Reverse 5’- AGA GGC CCG AGT AGG TGT AG) was performed using iTaq Universal SYBR Green Supermix (Bio-Rad laboratories). RefFinder [27] was used to establish the stability of the reference genes, and based on this and similar reaction efficiency to the target genes, cyclophilin (Forward 5’- TCT GCA CTG CCA AGA CTG AG, Reverse 5’- GTC CAC AGT CGG AGA TGG TG), B2M (Forward 5’- TGC TGT CTC CAT GTT TGA TGT ATC T Reverse 5’-TCT CTG CTC CCC ACC TCT AAG T) and ACTB (Forward 5’- CGA TAT CGC TGC GCT CGT, Reverse 5’- ATA CCC ACC ATC ACA CCC TG) were used as reference genes. qPCR was performed with a QuantStudio 7 Flex (Applied Biosystems, Foster City, CA). Primers were either adapted from existing literature or designed using Primer-BLAST (http://www.ncbi.nlm.nih.gov/tools/primer-blast/) to include all splice variants, and were purchased from Sigma-Aldrich. Primer specificity was confirmed from melting curve analysis. The PCR reaction contained 0.3 μM of each forward and reverse primer. A serial dilution analysis was used to determine the amount of template cDNA. The standard thermocycling program consisted of a 95°C denaturation pre-treatment for 10 min, followed by 40 cycles of 95°C for 15 s and 60°C for 60 s. All samples were run in duplicate with template free controls, and the mean Ct values were calculated. ΔCt was calculated as the difference between the target gene and the three reference genes. ΔΔCt was obtained by normalizing the ΔCt values of the treatments to the ΔCt values of Normal pH control at 0 h.

### Bioenergetics and mitochondrial respiration analyses

L6 myotubes were treated with normal, high and low pH media, with and without lactate, for five hours. They were then returned to normal media for 16 h before measurements of the bioenergetics profile of the cells were taken using the Seahorse XF24 Flux Analyser (Seahorse Bioscience). On the day of the measurements cells were washed and media replace with unbuffered DMEM (25 mM glucose, 1 mM pyruvate, 1 mM glutamate). Cells were incubated at 37 °C in a non-CO_2_ incubator for 1 h prior to bioenergetics assessment to allow the cells to adjust metabolism to 25 mM glucose. Three basal oxygen consumption rate (OCR) measurements were performed using the Seahorse analyser and measurements were repeated following injection of 1 μM oligomycin, 1 μM FCCP, 1 μM rotenone and 1 μM antimycin A. Respiratory parameters of mitochondrial function were calculated as described previously [28].

### Statistical analysis

All values are expressed as mean ± SEM. All protein content and gene expression results were normalized to the 0 h Normal pH sample. Data were analysed for statistical significance using the Univariate Analysis of Variance test, except for comparisons of protein phosphorylation or mRNA content between treatments where the students t-test was used. SPSS Statistics 22 was used for all statistical analysis. Significance was set at p ≤ 0.05.

## Results

### pH, and glucose and lactate concentrations, in incubation media

The media pH was significantly higher in the high pH manipulation groups and significantly lower in the low pH manipulations, when compared with the 0 h normal pH condition, when either the low or high glucose media was used (Figure 1a, 1b). The intracellular pH showed a similar pattern (Figure 1c, 1d). Lactate concentration remained constant throughout the incubations (Low glucose normal lactate 0.1 ± 0.0 to 1.3 ± 0.1, 20 mM Sodium lactate 15.7 ± 0.5 to 17.8 ± 0.3 mM, and High glucose normal lactate 0.3 ± 0.0 to 1.0 ± 0.1, 20 mM Sodium Lactate 17.5 ± 0.3 to 18.3 ± 0.4 mM) (Figure 1e, 1f). Glucose concentrations remained consistent between groups, with the exception of the low pH manipulations in both the low and high glucose media that did not see a drop in glucose concentration at 6 h (Low glucose media: Normal pH 4.8 ± 0.1, low pH 5.3 ± 0.1, low pH + lactate 5.3 ± 0.1 mM) (High glucose Normal pH 24.6 ± 0.3, low pH 25.2 ± 0.2, low pH + lactate 25.9 ± 0.3 mM) (Figure 1g, 1h).

### Cell viability

Measurements of cell viability, using trypan blue staining and a commercial cytotoxicity assay, showed that increasing the pH of the cell culture media to 8.1 or decreasing it to 7.0 does not result in significant changes in cell viability in either the low or high glucose media. The addition of 20 mM sodium L-lactate to the cell culture media also did not negatively affect cell viability, although in the high glucose media trypan blue staining did indicate an increase in cell viability with the addition of 20 mM sodium lactate (Figure 2).

**Figure 2:**
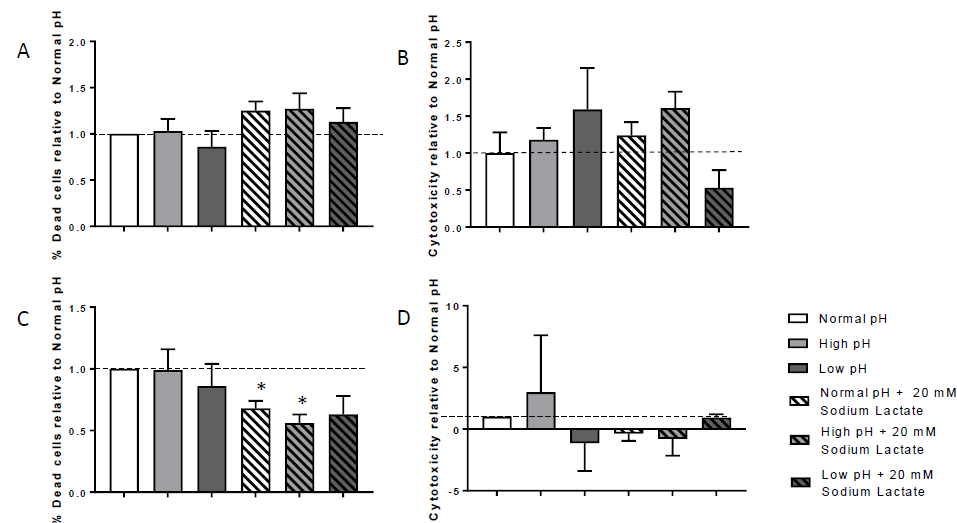
Cell viability after incubation of L6 myocytes in low or high glucose media. A. Trypan blue staining in low glucose media cells B. Low glucose media cytotoxicity C. Trypan blue staining in high glucose media cells D. High glucose media cytotoxicity * Significantly different from Normal pH group at the corresponding time, P ≤ 0.05. Values are means ± SEM. Samples are from 4 individual experiments.

### Effects of altered pH and lactate concentration on protein phosphorylation and localization Low glucose media

In conditions where the pH and lactate concentration were similar to that seen following high-intensity exercise (i.e., low pH and higher lactate concentration) [29], Akt (Ser473) phosphorylation was decreased at 1 h compared to Normal pH, whilst AMPK (T172) did not change significantly in the low glucose media (Figure 3a). However, low pH alone decreased AMPK phosphorylation at 1 h (Figure 3b). After 6 h, Akt and AMPK phosphorylation had returned to normal (Figure 3a,b) and the nuclear relative abundance of HDAC5 was decreased in both low pH conditions (Figure 4a).

**Figure 3:**
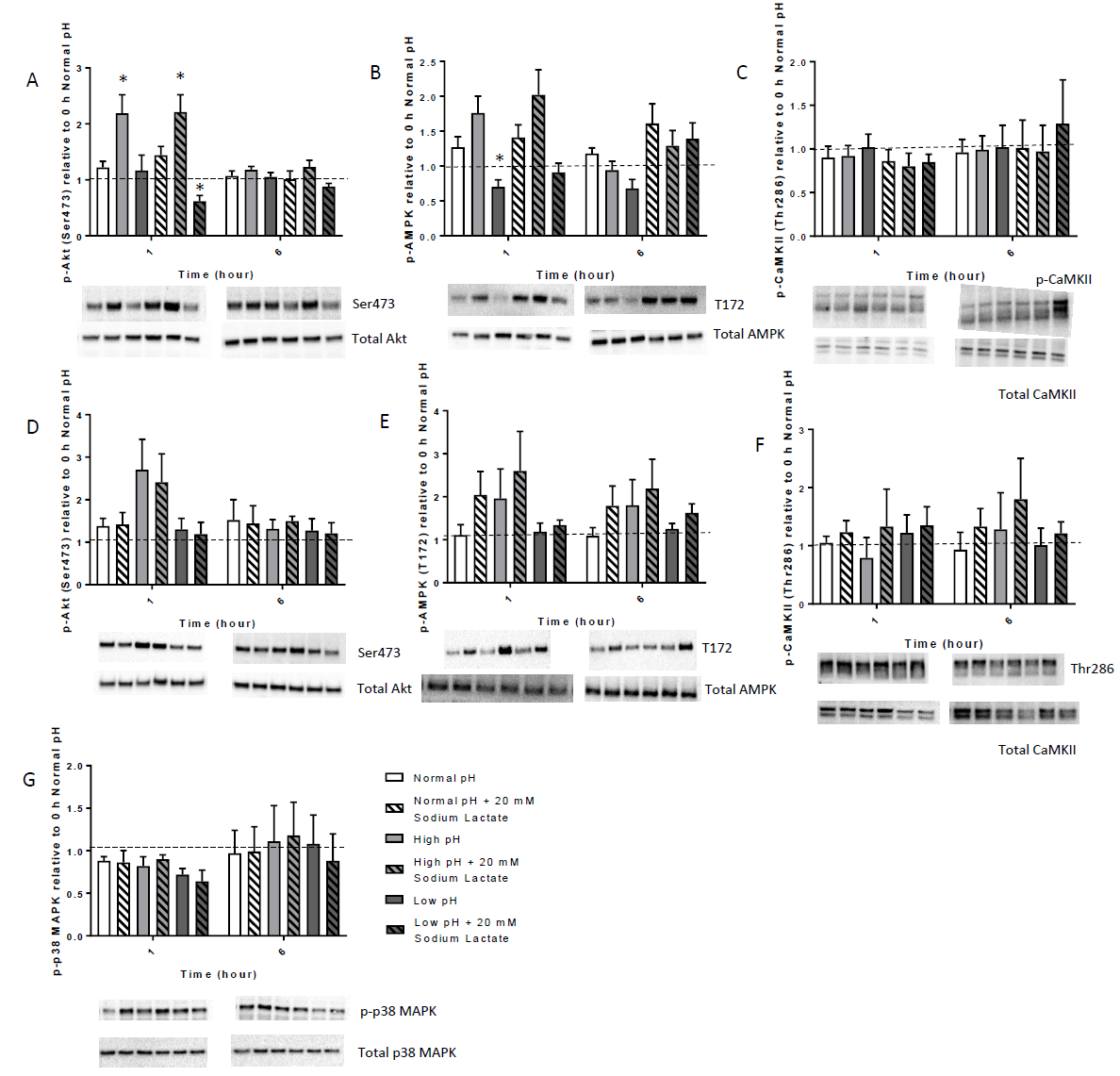
Protein phosphorylation A. Akt (Ser473) in low glucose media B. AMPKα (T172) in low glucose media. C. CaMKII (Thr286) in low glucose media D. Akt (Ser473) in high glucose media E. AMPKα (T172) in high glucose media. F. CaMKII (Thr286) in high glucose media G. p38 MAPK (residue) in high glucose media * Significantly different from the Normal pH group at the corresponding time P = ≤ 0.05. Values are means ± SEM. Samples are from five independent experiments.

**Figure 4:**
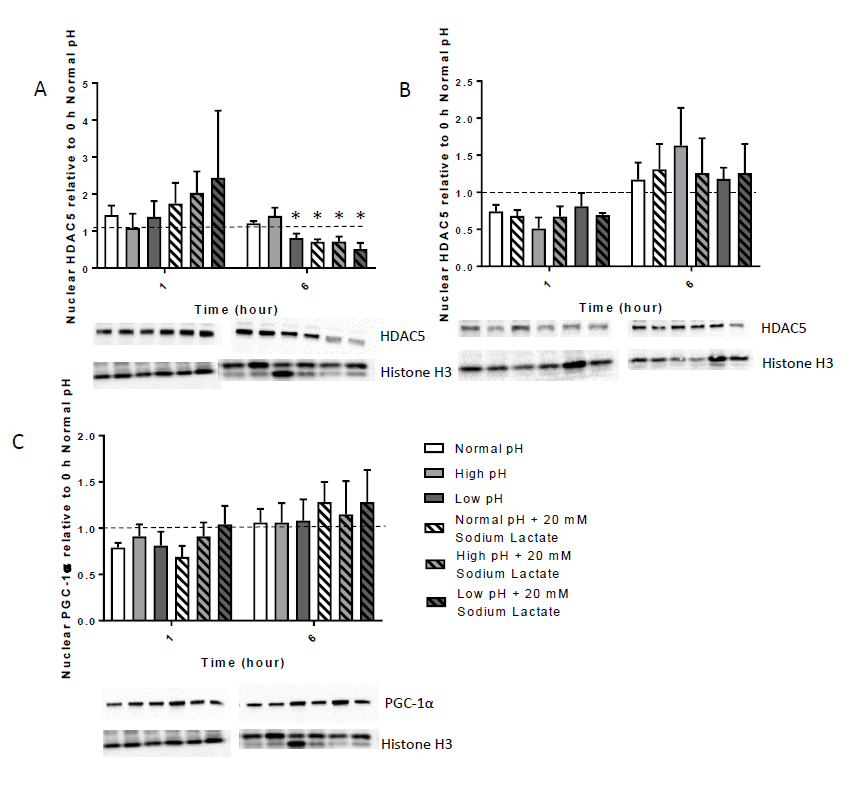
Nuclear localization. A. Low glucose HDAC5 nuclear localization. B. High glucose HDAC5 nuclear localization. C. Low glucose nuclear PGC-1α localization. * Significantly different from Normal pH group at the nominated time P = ≤ 0.05. Values are means ± SEM Samples are from four independent experiments.

When the pH was increased there was greater Akt (Ser473) phosphorylation (Figure 3a), whilst nuclear HDAC5 relative abundance was unchanged (Figure 4). CaMKII phosphorylation at Thr286 was not altered with any of the treatments (Figure 3c and f), nor was p38 MAPK phosphorylation (Figure 3). In all three 20 mM lactate conditions there was decreased nuclear localization of HDAC5 at 6 h, but not at 1 h (Figure 4a). PGC-1α nuclear localization was not altered significantly with any treatment (Figure 4c).

### High glucose media

In cells incubated in high glucose media none of the manipulations resulted in any significant changes in protein phosphorylation or protein localization of the proteins studied (Figures 3 and 4).

### Gene expression

Less HDAC5 in the nucleus is linked with de-repression of gene transcription [30], which is consistent with previous research reporting that increased lactate can increase the transcription of MCT1, basigin (also known as CD147), and PGC-1α [13]. Therefore, we then looked at the mRNA content of a genes encoding transcription factors or proteins with a role in lactate transport or mitochondrial biogenesis.

### No change in the expression of genes encoding proteins important for lactate transport

The mRNA content of MCT1 was not changed with any of the treatments (Figure 5a and c). There were also no significant change in CD147 (basigin) mRNA content (Figure 5b and d), with the exception of a significant decrease at 1 h in the high glucose media with high pH and lactate.

**Figure 5:**
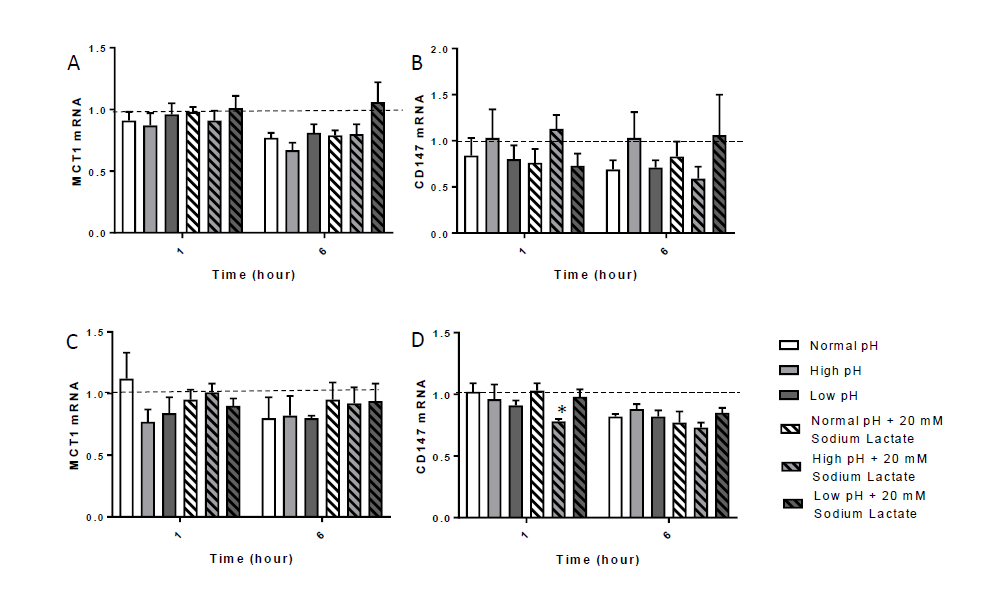
mRNA expression of A. MCT1 and B. CD147 when cells were incubated in low glucose media and mRNA expression of C. MCT1 and D. CD147 when cells were incubated in high glucose media. Values are means ± SEM and expressed relative to 0 h Normal pH. There were no significant differences between conditions. Samples are from 4 to 5 independent experiments.

### Genes implicated in the activation of mitochondrial biogenesis

There were no significant changes in the mRNA content of NRF1, NRF2, Tfam, COXIV, or cytochrome c with either high or low pH or an increased media lactate concentration (Figure 6). PGC-1α mRNA content was not changed after one hour of altered pH or lactate; however, a 6-h exposure to a high pH significantly decreased PGC-1α expression by approximately 40% with and without additional lactate. This effect was consistent in both the low and high glucose media (Figure 7a and d). The mRNA content of splice isoforms PGC-1α1 and PGC-1α4 was not significantly altered in most conditions and at most time points (Figure 7b – f). However, there was a significant decrease in PGC-1α4 expression with high pH and lactate in the high glucose media, but not the low glucose media (p = 0.058) (Figure 7f).

**Figure 6:**
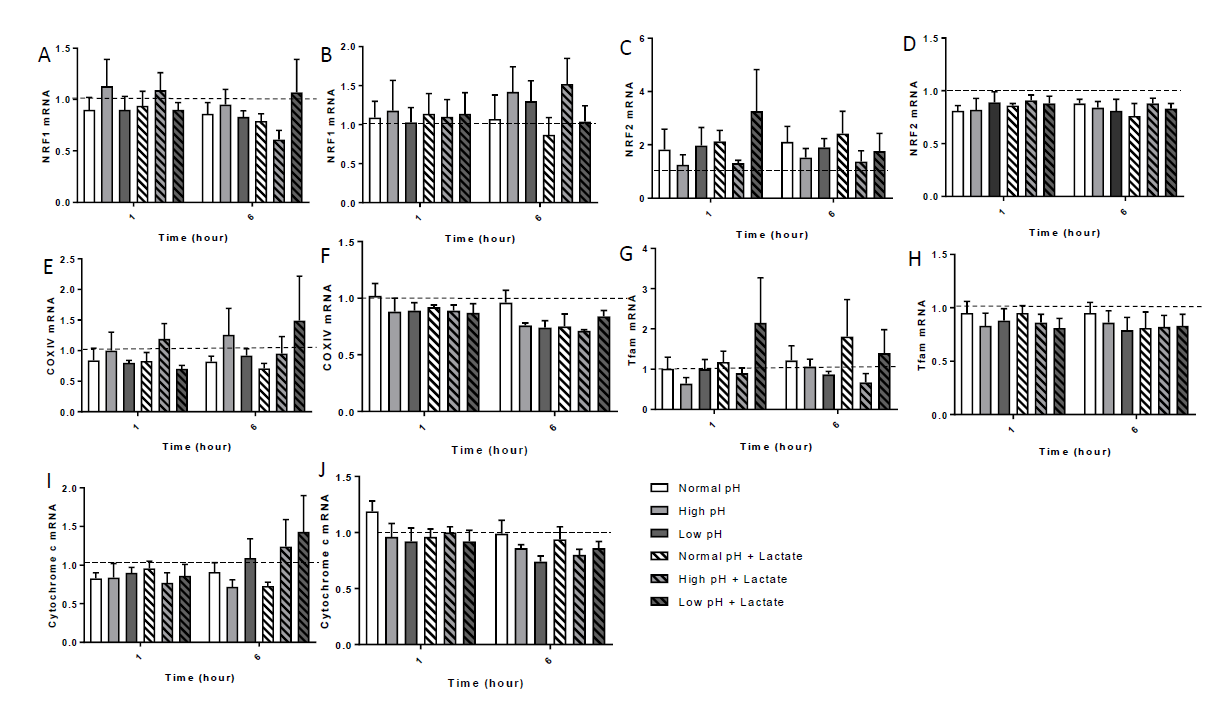
mRNA expression of mitochondrial genes in cells treated with low or high glucose media. A. Low glucose NRF1, B. High glucose NRF1 C. Low glucose NRF2, D. High glucose NRF2, E. Low glucose COXIV, F. High glucose COXIV, G. Low glucose Tfam, H. High glucose Tfam, I. Low glucose cytochrome c, J. High glucose cytochrome c.* Significantly different from normal pH group at nominated time P = ≤ 0.05. Values are means ± SEM relative to 0 h Normal pH. Samples are from five independent experiments.

**Figure 7:**
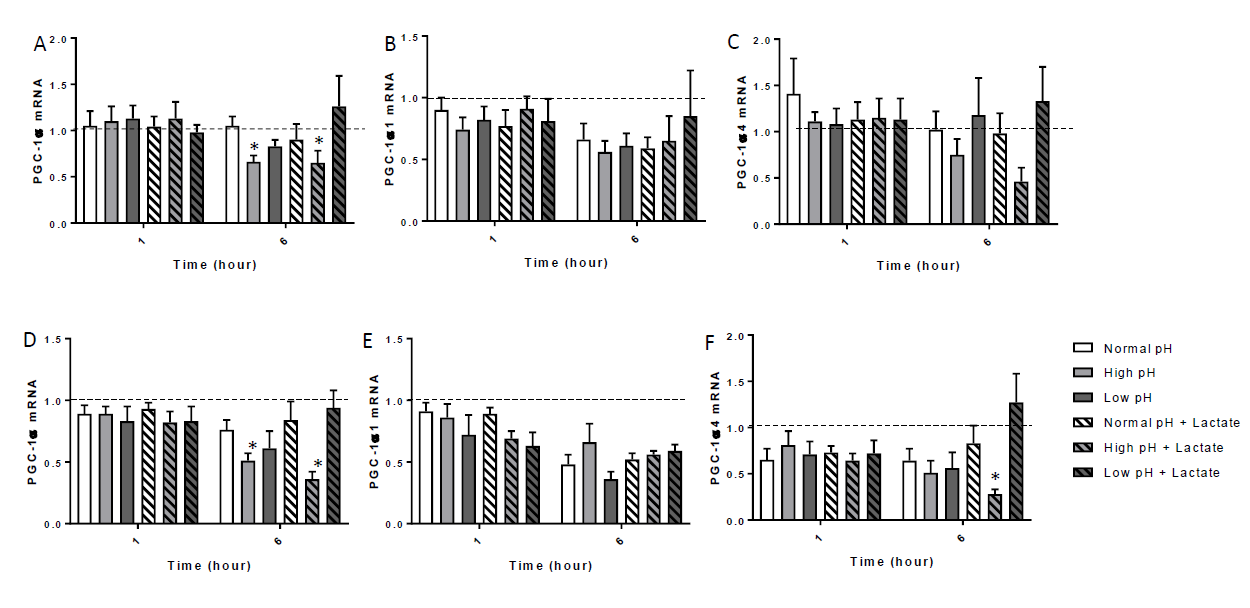
mRNA expression of Peroxisome proliferator-activated receptor gamma coactivator 1-alpha (PGC-1α) and two of it’s isoforms in cells treated with low or high glucose media A. Low glucose PGC-1α, B. Low glucose PGC-1α1, C. Low glucose PGC-1α4, D. High glucose PGC-1α, E. High glucose PGC-1α1, F. High glucose PGC-1α4 * Significantly different from normal pH group at corresponding time. P = ≤ 0.05. Values are mean ± SEM relative to 0 h normal pH. Samples are from four independent experiments.

### Bioenergetics and mitochondrial respiration analyses

Exposure to both high and low pH media significantly decreased basal mitochondrial respiration, ATP turnover, and maximum mitochondrial respiratory capacity. There was no effect of lactate alone on mitochondrial respiration; however, addition of lactate to the ‘high’ and ‘low’ media appeared to return mitochondrial function to normal or at least blunt the effects of the high or low pH (Figure 8).

**Figure 8:**
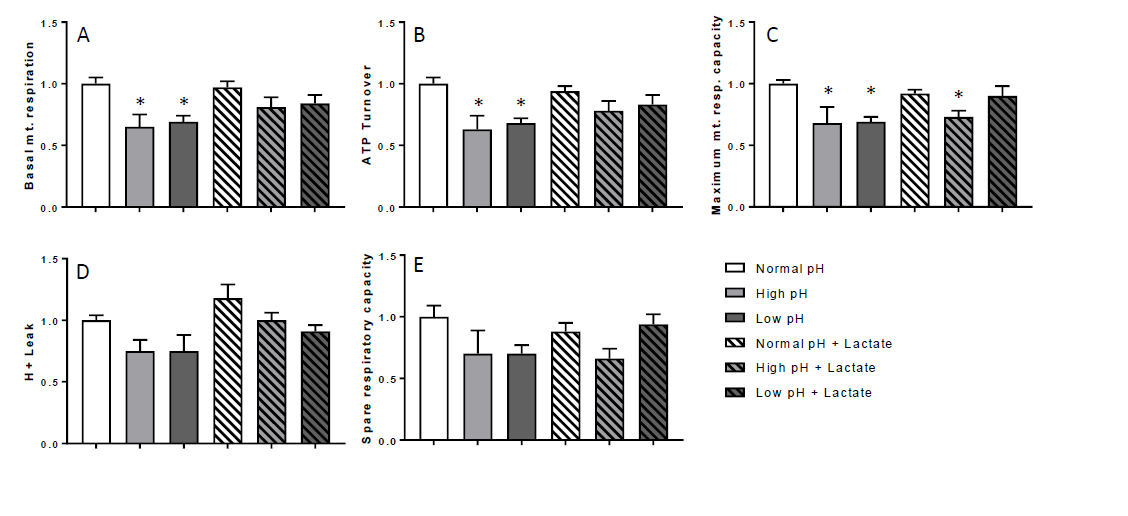
The effects of manipulating cell media pH and lactate concentration on mitochondrial function in L6 myocytes. A. Basal mitochondrial respiration B. ATP turnover C. Maximum mitochondrial respiratory capacity D. H^+^ leak E. Spare respiratory capacity * Significantly different from normal pH group P = <0.05. Values are mean ± SEM.

## Discussion

This is the first study to examine the impact of a low, normal, or high pH, with and without high physiological concentrations of lactate, on markers of mitochondrial biogenesis and function in L6 myocytes. In general, there were few significant effects of these manipulations. However, a low pH (approximately 6.8, as seen immediately after high-intensity exercise) decreased p-Akt and p-AMPK relative abundance in the cytoplasm and also decreased HDAC5 relative abundance in the nucleus. Increasing intracellular pH increased Akt phosphorylation, but without an effect on HDAC5 relative abundance in the nucleus unless it was also accompanied by an increase in lactate concentration. Increasing media pH also decreased the expression of PGC-1α mRNA at 6 h. The most consistent finding was that increasing the lactate concentration for 6 h decreased the relative abundance of HDAC5 in the nucleus. Mitochondrial respiration was decreased with both a low and high media pH.

In this study we examined the response of genes and proteins known to have a role in mitochondrial biogenesis to physiologically-relevant changes in pH and lactate [14], which did not negatively affect cell viability. While greater, non-physiological changes may have produced different results, greater changes have also been reported to negatively affect cell viability [31]. By changing the media pH we were able to also alter the intracellular pH (Figure 1). As expected, due to the buffering capacity of cells [32], the alteration in intracellular pH was not as great as the changes in extracellular pH. To enable comparison with previous literature, we completed two sets of experiments; we performed one manipulation in low and one manipulation in high glucose containing media. We observed that most of the significant changes occurred in low glucose αMEM incubated cells, which is most similar to blood glucose levels *in vivo*.

A decrease in pH was associated with a decrease in p-Akt relative abundance, but only when accompanied by an increase in lactate concentration (as occurs during muscle contraction). In contrast p-Akt relative abundance increased with an increased pH (with or without the addition of 20 mM Lactate). Metabolic acidosis, in an animal model of chronic kidney disease, has previously been reported to be associated with a decrease in p-Akt content [17]. However, another study in human carcinoma cells and immortalized fibroblasts found that acidification of the cell culture medium from 7.4 to 6.4 did not affect phosphorylation of Akt [33]. Previous reports have also shown that the Akt and MAPK pathways interplay at different levels and that they may be part of a negative feedback loop [34]. However, despite the observed changes in p-Akt relative abundance we did not see changes in p-p38 MAPK content in the current study. Thus, the implications of a decrease or increase in p-Akt in response to a change in pH are unclear. However, given the role of Akt in muscle synthesis and metabolism [35], a decrease in pH may have a negative effect on metabolism and muscle synthesis. This is reflected by the higher glucose concentration in the media after 6 h incubation in low pH media in this study.

Another important signalling protein, activated in response to stress (such as exercise), is AMPK [36]. In the present study, there was a decrease in p-AMPK relative abundance in the low pH with lactate condition, as well as a trend towards increased phosphorylation with a higher pH. Consistent with our study, Zhao et al [31] observed that an acidic pH decreased p-AMPK relative abundance, whilst an alkaline pH increased p-AMPK relative abundance in cultured cardiomyocytes [31]. Another study in cultured fibroblasts also found that an acidic or low pH decreased p-AMPK relative abundance [33]. An increase in p-AMPK relative abundance has been linked to increased mRNA content of proteins favouring oxidative phosphorylation, such as PGC-1α and cytochrome c [7, 36-38]. This inducement of mitochondrial biogenesis by p-AMPK is thought to occur by alteration of the binding activity of transcription factors, such as NRF1 and MEF2, as well as altered localization of HDACs [2, 37, 39]. Therefore, a decrease in AMPK phosphorylation suggests a potential for decreased mitochondrial biogenesis with a lowered pH.

As AMPK phosphorylation has been reported to affect HDAC localisation [40], we next examined the nuclear localisation of HDAC5. We observed a decrease in HDAC5 nuclear relative abundance after a 6 h incubation with additional lactate and/or a low pH. Less nuclear HDAC5 suggests an increased opportunity for gene transcription [41]. Thus, the decrease in nuclear HDAC content after the addition of lactate is consistent with the increased transcription of PGC-1α reported in a similar, previous study [13]. In contrast, we did not observe any significant increases in gene transcription in the present study (with the exception of a decrease in PGC-1α mRNA content with an increased pH), despite using an identical lactate concentration and the same cell line. It is difficult to explain these contrasting findings, but we note that the changes reported by Hashimoto et al [13] were small. Additionally, as we did not observe a decrease in HDAC5 nuclear protein abundance until 6 h, it may be that greater time is required for this change in HDAC5 to promote significant increases in gene transcription.

In addition to the changes we saw in protein phosphorylation and localization, and minor changes in mRNA content, altering the media pH above or below its normal range decreased mitochondrial function (as measured by parameters such as basal mitochondrial respiration, ATP turnover and maximum mitochondrial respiratory capacity). To account for the time effects of mitochondrial adaptations, these measurements were undertaken 16 h after the exposure to the altered pH medium; our results suggest that alterations in extracellular pH may either have prolonged effects beyond the time of actual pH change or that changes in mitochondrial respiration (and associated signaling events) may not occur immediately upon a pH change but at later time points. Therefore, it may be useful for future research to also examine protein phosphorylation and expression changes at time points beyond those measured in this study.

In conclusion, we observed that short-term physiological alterations in extracellular pH and lactate result in alterations in AMPK and Akt phosphorylation and HDAC5 localization, suggesting the potential for alterations in mitochondrial biogenesis and function. Indeed, we found that mitochondrial function was decreased with both a high and low pH. This was accompanied by changes in the mRNA expression of PGC-1α with a high pH. However, we did not observe any alterations in the expression or activation of a number of other proteins or genes proposed to be involved in mitochondrial biogenesis. Due to the transient nature of changes in mRNA expression and protein activation, it is possible we were not able to detect some changes that may have occurred. Future work will be required to establish if changes in mRNA expression occur at time points beyond 6 h.

## Funding

This study was supported by a grant from the Australian Research Council (ARC) to DJB.

---

**Table 1:** Summary of results

| **pH** | **Normal** |  | **High** |  | **Low** |  |
| --- | --- | --- | --- | --- | --- | --- |
| **Lactate** | - | + | - | + | - | + |
| **Protein Signalling** |  |  |  |  |  |  |
| p-Akt | - | - | ↑ | ↑ | - | ↓ |
| p-AMPK | - | - | - | - | ↓ | - |
| p-CaMKII | - | - | - | - | - | - |
| Nuclear PGC-1α | - | - | - | - | - | - |
| Nuclear HDAC5 | - | ↓ | - | ↓ | ↓ | ↓ |
| **Gene Expression** |  |  |  |  |  |  |
| PGC-1α | - | - | ↓ | ↓ | - | - |
| **Mitochondrial Respiration** |  |  |  |  |  |  |
| Basal mt. resp. | - | - | ↓ | - | ↓ | - |
| ATP turnover | - | - | ↓ | - | ↓ | - |
| Max. mt. resp. capacity | - | - | ↓ | ↓ | ↓ | - |

Abbreviations: AMP-activated protein kinase (AMPK), Ca<sup>2+</sup>/calmodulin-dependent protein kinase II (CaMKII), Peroxisome proliferator-activated receptor gamma coactivator 1-alpha (PGC-1α), histone deacetylase 5 (HDAC5), basal mitochondrial respiration (Basal mt. resp.),

## Abbreviations

(AMPK): AMP-activated protein kinase
(CaMKII): Ca2+/calmodulin-dependent protein kinase II
(PGC-1α): Peroxisome proliferator-activated receptor gamma coactivator 1-alpha
(HDAC5): histone deacetylase 5
(Basal mt. resp.): basal mitochondrial respiration
(Max. mt. resp. capacity): maximum mitochondrial respiratory capacity
‘p’: prefix refers to phosphorylation

